# Cerebral organoids for modeling of HSV-1-induced-multiscale neuropathology associated with Alzheimer’s disease and phenotypic rescue

**DOI:** 10.1101/2022.02.11.480178

**Authors:** Haowen Qiao, Wen Zhao, Moujian Guo, Lili Zhu, Tao Chen, Jibo Wang, Zhentao Zhang, Ying Wu, Pu Chen

## Abstract

Herpes simplex virus type I (HSV-1) infection is a potential risk factor involved in the pathogenesis of Alzheimer’s disease (AD) and contributes to the AD viral hypothesis. However, further understanding of the neuropathological effects of the HSV-1 infection is hampered by the limitations of existing infection models due to the distinct differences between human brains and other mammalians’ brains. Here we generated cerebral organoid models derived from pluripotent stem cells to investigate the HSV-induced pathogenesis associated with AD and the role of antiviral drugs in the phenotypic rescue. Our results identified that the HSV-1-infected cerebral organoids recapitulated multiscale neuropathology pertaining to the onset and progression of AD across the genetic, cellular, and tissue levels, including the multicellular Aβ deposition, dysregulated endogenous AD mediators, reactive gliosis, neuroinflammation, and neural loss. Furthermore, we identified that two antiviral drugs, namely Ribavirin (RBV) and Valacyclovir (VCV), inhibited HSV-1 replication and rescued the neuropathological phenotypes associated with AD in the HSV-1-infected cerebral organoids, implying their therapeutic potential to slow down the progression of AD. Our study provides a high-fidelity human-relevant in-vitro HSV-1 infection model to reconstitute the multiscale neuropathological features associated with AD and discover therapeutic drug candidates relevant to the AD viral hypothesis.

**IMPORTANCE:** AD is a progressive neurodegenerative disorder characterized by neuronal death, brain atrophy, and cognitive impairment. HSV-1 is a neurotropic double-stranded DNA virus resulting in long-term neurological sequelae in adults.

There is accumulating evidence indicating that AD may be associated with HSV-1 infections. In this study, we generated human-physiologically related cerebral organoids derived from human embryonic stem cells (hESCs) to investigate the influence of HSV-1 infection on the neuropathology associated with AD. We found that the HSV-1-infected cerebral organoids could model typical neuropathological features associated with AD. Additionally, we discovered that both RBV and VCV significantly rescued the HSV-1-induced pathological phenotypes associated with AD. These findings may open new avenues of consideration related to HSV-1 infection and AD. Furthermore, the contribution of these antiviral drugs to AD linked to HSV-1 infection will provide new clues to potential new therapies.

## INTRODUCTION

Alzheimer’s disease (AD) is a progressive neurodegenerative disorder characterized by neuronal death, brain atrophy, cognitive impairment, and ultimately a decline in daily life activities.^1^ The cost of AD healthcare is $818 billion in 2015 worldwide and could rise as high as $2 trillion by 2030.^2^ Approximately 50 million individuals around the world suffer from AD, and the number of AD patients is predicted to be 152 million in 2050 according to the World Alzheimer Report^3^. Therefore, there is an urgent need for probing AD pathogenic mechanisms and develops effective therapeutics.

Clinically, AD is classified into sporadic AD (SAD) and familial AD (FAD) based on the age of onset. More than 95% of AD cases are SAD.^4^ The common neuropathological hallmarks of SAD and FAD are mainly characterized by the formation of intracellular neurofibrillary tangles by hyperphosphorylated tau proteins, and the extracellular sediment of insoluble aggregates amyloid-β protein (Aβ) in the cortex and subcortex. There exists various hypotheses regarding the SAD etiology, including the amyloid hypothesis, tau propagation hypothesis, cholinergic hypothesis, inflammatory hypothesis, viral hypothesis, and so on.^5^ Notably, some recent studies have provided intriguing implications that the AD viral hypothesis and the amyloid hypothesis could be complementary with each other and provide an intrinsic and consistent interpretation that the infection can trigger the Aβ production, ultimately resulting in the onset of AD pathology. However, the ultimate etiology of AD remains controversial.

In the resent years, there is increasingly accumulating evidence indicating that AD may be associated with various pathogens, such as herpes simplex virus type I (HSV-1) and cytomegalovirus.^6^ Especially, HSV-1 is highly prevalent and capable of establishing lifelong infection, which accounts for a heavy disease burden in both life quality and economy.^7^ HSV-1 is a neurotropic double-stranded DNA virus that transmits across the blood-brain barrier (BBB) in the central nervous system, resulting in long-term neurological sequelae in adults. The clinical studies^8^ have reported a molecular, genetic, and clinical network analysis of virome from the human post-mortem brains and concluded that levels of human herpesviruses were significantly higher in the subjects with AD pathology than the subjects in the controls. Notably, a retrospective cohort study,^9^ following 33,448 people in Taiwan, suggested that the people who experienced herpesvirus infections had a 2.5-fold increased risk in developing AD compared to the people without the infections. All these clinical evidences imply that the AD pathogenesis is associated with the HSV-1 infection. Thus, there is a further demand for revealing the causal relationship between the HSV-1 infection and the onset and progression of AD pathology.

Wild-type mice don’t intrinsically develop AD-associated neuropathological phenotypes. Thus, the study of the AD pathogenesis currently relies on the transgenic mouse models that overexpress pivotal risk genes of FAD.^10^ These transgenic mouse models could recapitulate some pathological features of AD, and be widely used in probing the underlying mechanisms of AD. For example, the 5×FAD transgenic mouse is a widely-recognized FAD mouse model that possesses a total of five typical familial mutations in amyloid precursor protein (APP) and presenilin 1 (PSEN1) genes discovered in the FAD patients. Using this model, Giovanna De Chiara and coworkers ^11^ found that HSV-1 infection accelerated Aβ deposition in the brain. The follow-up studies^12, 13^ further revealed that Aβ oligomers aggregated around the herpesvirus particles in the cortex, implying that Aβ oligomers may act as a type of endogenous antiviral protein in the innate immune system. However, the transgenic mouse models could not faithfully mimic the pathogenesis of SAD that do not involved a genetic background of familial mutations. Besides, there are distinct neurophysiological differences between the human and the mouse brains across the multiple bio-hierarchical levels at the varied neurodevelopmental stages, including morphogenesis of cortex, cytoarchitecture of cortex, neuro polarization of neuroepithelium, and self-organization of cerebral cortex. Therefore, the transgenic mice are not faithful research models to study the human-specific neuropathophysiological characteristics in the late-on AD patients.

Development of human physiologically relevant infection models are vital to understand how virus infections impact the onset and progression of AD. Fortunately, emerging cerebral organoids offer a more human relevant in-vitro model to study the neurophysiology and disease of human cerebral cortex.^14^ Specifically, cerebral organoids could emulate the human cerebral cortex’s cellular diversity,^15^ such as the intermediate progenitors and the neurogenic outer radial glia. Furthermore, the cerebral organoids can resemble the specific cytoarchitecture in the human cerebral cortex, including the cortical lumen organization^16^ and cortical layers^17^. Recently, the cerebral organoids have been used to model neurodegenerative diseases^18^ and neurotropic virus infections^19^. Cesar Gonzalez and his colleagues established cerebral organoid models derived from FAD patients’ induced pluripotent stem cells, and these models recapitulated some typical neuropathological features in the AD-patient brains, including Aβ accumulation, p-tau aggregation, and cellular apoptosis.^20^ Moreover, in our previous study, we generated the HSV-1-infected cerebral organoids to reconstitute the neuropathological features of the neurodevelopmental disorders in the human fetal brain.^21^ Besides, Leonardo D’Aiuto et al. used cerebral organoids to study the acute and latent HSV-1 infection, providing substantial evidence that the cerebral organoids are suitable to model HSV-1-CNS interactions.^22^

In this study, we generated human-physiologically related cerebral organoids derived from human embryonic stem cells (hESCs) to investigate the influence of HSV-1 infection on the neuropathology associated with AD. We found that the HSV-1-infected cerebral organoids could model typical neuropathological features associated with AD including multicellular Aβ deposition and dysregulated exogenous AD mediators. Notably, the HSV-1 infected cerebral organoids also demonstrated reactive gliosis, neuroinflammation, and neuron loss. Furthermore, we discovered that both Ribavirin (RBV) and Valaciclovir (VCV) significantly suppressed viral replication and rescued the HSV-1-induced multiscale pathological phenotypes associated with AD, implying their therapeutic effects on human AD patients. Taken together, we developed effective human brain infection models for mimicking the HSV-1-induced multiscale neuropathological phenotypes and further supported the AD viral hypothesis. We expected this model will facilitate the applications to explore therapeutic intervention and target HSV viral reservoirs relevant to AD.

## RESULTS

### HSV-1 led to AD-associated Aβ deposition in the cerebral organoids

To investigate the susceptibility of human brain cells to HSV-1 infection, we tested hESC-derived cerebral organoids generated via the previous protocols. [19] The cerebral organoids at the different growth stages (day14, D14; day 42, D42; day 65, D65) were inoculated with HSV-1 for three days (D14+3, D42+3, D65+3), then we examined the cerebral organoids after inoculum removal (Fig. 1A and 1B). The quantitative 3D immunofluorescence imaging of cerebral organoids revealed that HSV-1 infection increased the Aβ expression at D14+3 (Fig. 2A), the results also showed that HSV-1 infection induced the Aβ deposition in at D42+3 (Fig. 2B) and D65+3 (Fig. 2C).

**Fig. 1.**
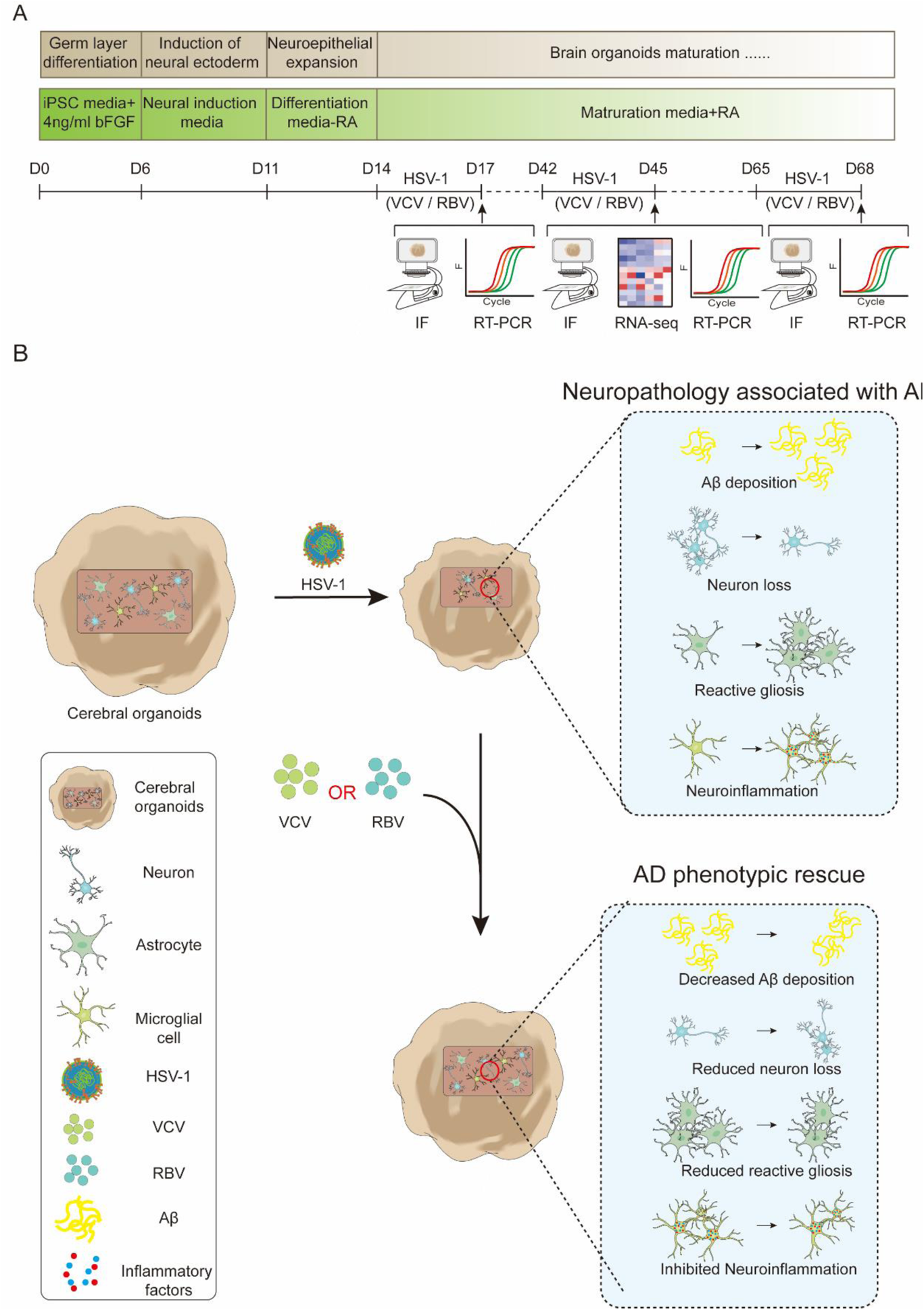
Schematic diagram of cerebral organoids for modeling of HSV-1-induced-multiscale neuropathology associated with Alzheimer’s disease and phenotypic rescue. (A) Schematic procedure of HSV-1-infected hESCs derived cerebral organoid timing and method. (B) Cerebral organoids for modeling of HSV-1-induced-multiscale neuropathology associated with Alzheimer’s disease and phenotypic rescue. HSV-1-infected cerebral organoids could model typical neuropathological features associated with AD including multicellular Aβ deposition, neuron loss, reactive gliosis, and neuroinflammation. Furthermore, both RBV and VCV significantly rescued the HSV-1-induced multiscale pathological phenotypes associated with AD.

**Fig. 2.**
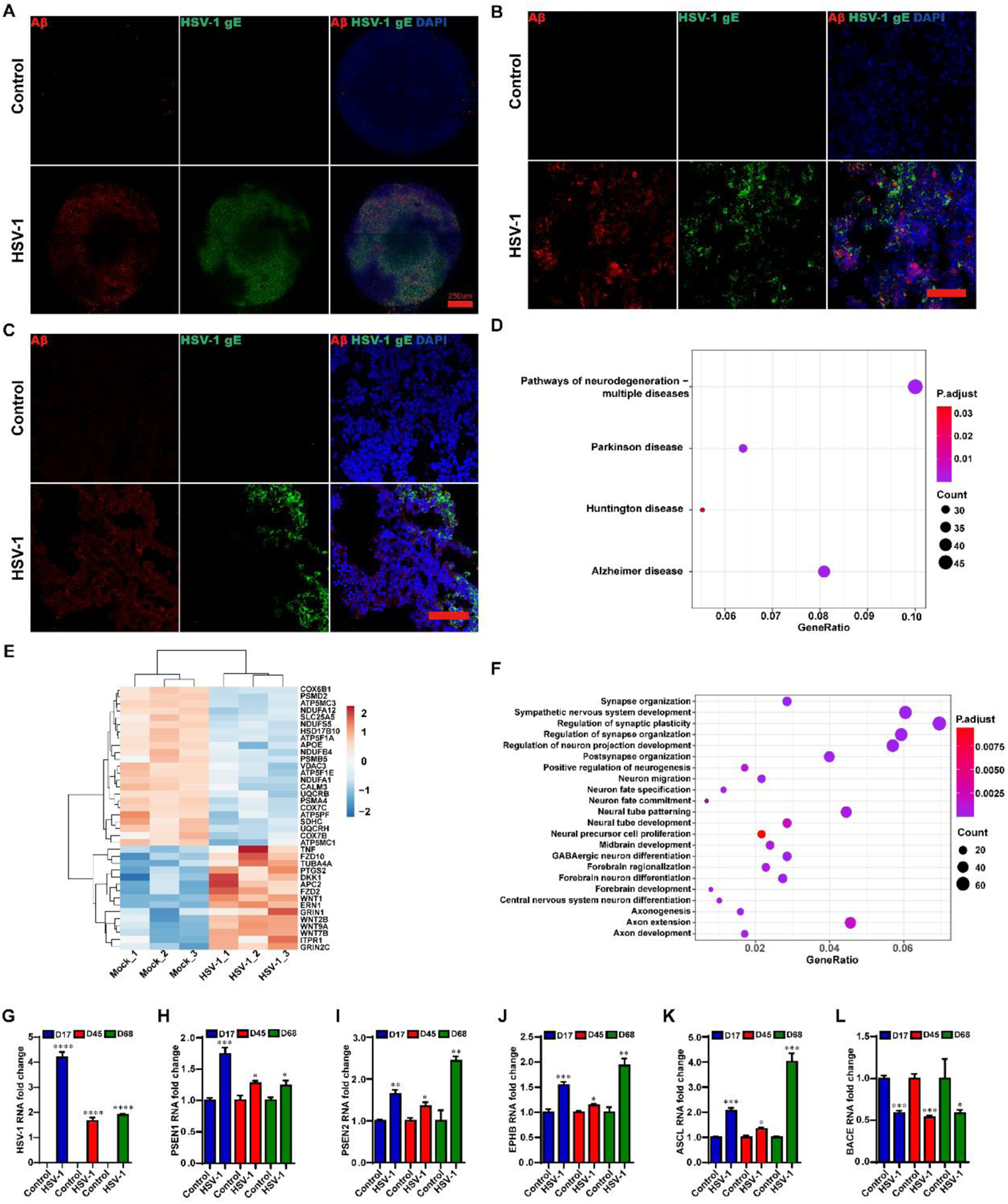
HSV-1 led to AD-associated Aβ deposition in the cerebral organoids. (A) Whole-mount 3D immunofluorescence imaging of HSV-1 and Aβ in the cerebral organoids with or without HSV-1 infection at D14+3. Scale bars: 250 µm. (B and C) The immunostaining for the HSV-positive and Aβ-positive cells in the cerebral organoids with or without HSV-1 infection at D42+3 and D65+3. Scale bars: 100 µm. (D and E) KEGG pathways enriched in the neurodegenerative diseases and the AD-related dysregulated genes in the cerebral organoids infected with the HSV-1, compared to the control. (F) Dot plot of selected enriched GO terms for the biological process for dysregulated genes in the cerebral organoids infected with the HSV-1, compared to the control. (G-L) RT-PCR analysis to monitor the mRNA expressions of HSV-1, APOE, PSEN1, PSEN2, ASCL, and BACE in the HSV-1 infected cerebral organoids at D14+3, D42+3 and D65+3. The expression value was normalized to the GAPDH expression level. Data represent the mean ± SEM. *p<0.05 vs the control group, **p<0.01 vs the control group, ***p<0.005 vs the control group, ****p<0.001 vs the control group, (n=6).

The RNA-seq analysis was also conducted to examine the regulatory effects of HSV-1 on the AD-associated gene networks in the cerebral organoids at D14+3. The KEGG pathway enrichment analysis (FDR < 0.05) indicated that the regulation of gene networks in the HSV-1-infected cerebral organoids were closely linked to neurodegenerative diseases, especially for AD (Fig. 2D). The heat map (Fig. 2E) further indicated that 30 AD-associated genes were highly responsive to HSV-1 infection, predicting that HSV-1 infection may play an essential regulatory role in the AD pathogenesis. The GO enrichment analysis of the differentially expressed genes (Fig. 2F) showed that the highly enriched categories were related to synapse organization, synapse assembly, neural tube development, nervous system development, axon guidance, cell morphogenesis neuron differentiation, neuron projection guidance, and neuron projection development. The remarkable convergence of KEGG pathway enrichment analysis and GO enrichment analysis prompted us to hypothesize that HSV-1 infection led to the AD-associated neuropathology in the cerebral organoids.

Then we determined the expression of widely recognized AD mediator genes, including PSEN1, PSEN2, EPHB, ASCL, and BACE in the cerebral organoids. The RT-PCR analysis indicated that HSV-1 infection caused up-regulation of HSV-1, PSEN1, PSEN2, EPHB, and ASCL at mRNA levels (Fig. 2G to 2K) and down-regulation of BACE mRNA expression in the cerebral organoids (Fig. 2L) at D14+3, D42+3, and D65+3, compared to the uninfected groups.

### HSV-1 infection resulted in the neuron loss

In order to test whether HSV-1 infection inhibit neuronal differentiation. We examined neuronal differentiation hallmarks of different stage in the infection models. The HSV-1 infected cerebral organoids exhibited a loss in the neuronal-associated protein (TUJ and MAP2) by immunofluorescence (Fig. 3A to 3C) at D65+3. In addition, the mRNA expressions of TUJ and MAP2 in HSV-1-infected cerebral organoids were also examined by real-time PCR at D42+3 and D65+3 (Fig. 3E to 3F), which were consistent with the results observed in immunofluorescence analysis, indicating a distinct neuron loss by HSV-1 infection.

**Fig. 3.**
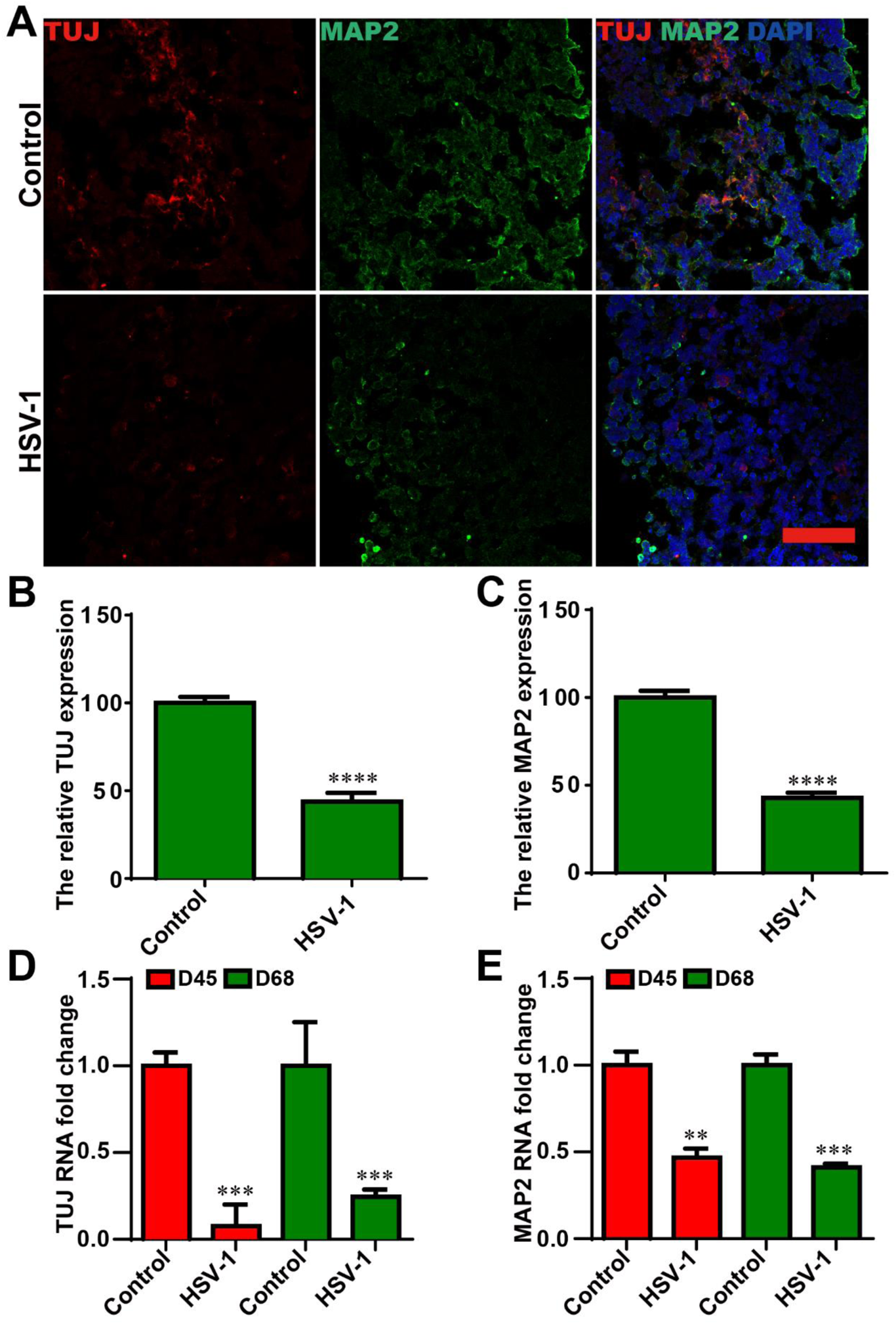
HSV-1 infection resulted in the neuron loss. (A) Co-immunostaining of TUJ and MAP2 were identified by immunofluorescence in the HSV-1 infected hESCs derived cerebral organoids at D65+3, Scale bars: 100µm. (B and C) The relative fluorescence intensity statistics of TUJ and MAP2 expressions were shown in different groups. (D and E) RT-PCR analysis to monitor the mRNA expressions of TUJ and MAP2 in HSV-1-infected cerebral organoids at D42+3 and D65+3. The expression value was normalized to the GAPDH expression level. Data represent the mean ± SEM. **p<0.01 vs the control group, ***p<0.005 vs the control group, ****p<0.001 vs the control group, (n=6).

### HSV-1 infection led to reactive gliosis and neuroinflammation in the cerebral organoids

It is widely accepted that glial-mediated reactive gliosis and neuroinflammation contribute to the progression of AD. Thus we examined the reactive gliosis and neuroinflammation in the HSV-1 infected cerebral organoids. CD11b is a cell surface integrin in the microglia, and CD68 is a marker of microglial lysosomes indicative of phagocytic microglia. The overexpression of CD11b and CD68 in active microglia is considered as the reactive gliosis in AD. We assessed the reactive gliosis in the organoids using immunofluorescence staining against these two markers at D65+3. The results indicated that the relative fluorescence intensity of CD11b and CD68 were significantly higher in the HSV-1-infected cerebral organoids compared to the control group (Fig. 4A to 4C). In addition, the expressions of GFAP (a marker of astrocytes) was highly up-regulated in response to HSV-1 infection by immunofluorescence staining (Fig. 4D and 4E). Similarly, the mRNA expressions of GFAP in HSV-1-infected cerebral organoids were also examined by real-time PCR at D42+3 and D65+3 (Fig. 4F), which were consistent with the results observed in immunofluorescence analysis What is more, the apparent decrease in mRNA expressions of the activated microglia markers (CD11b, CD68, CXC3R, and HLADR) were observed in the cerebral organoids at D42+3 and D65+3 (Fig. 4G to 4J).

**Fig. 4.**
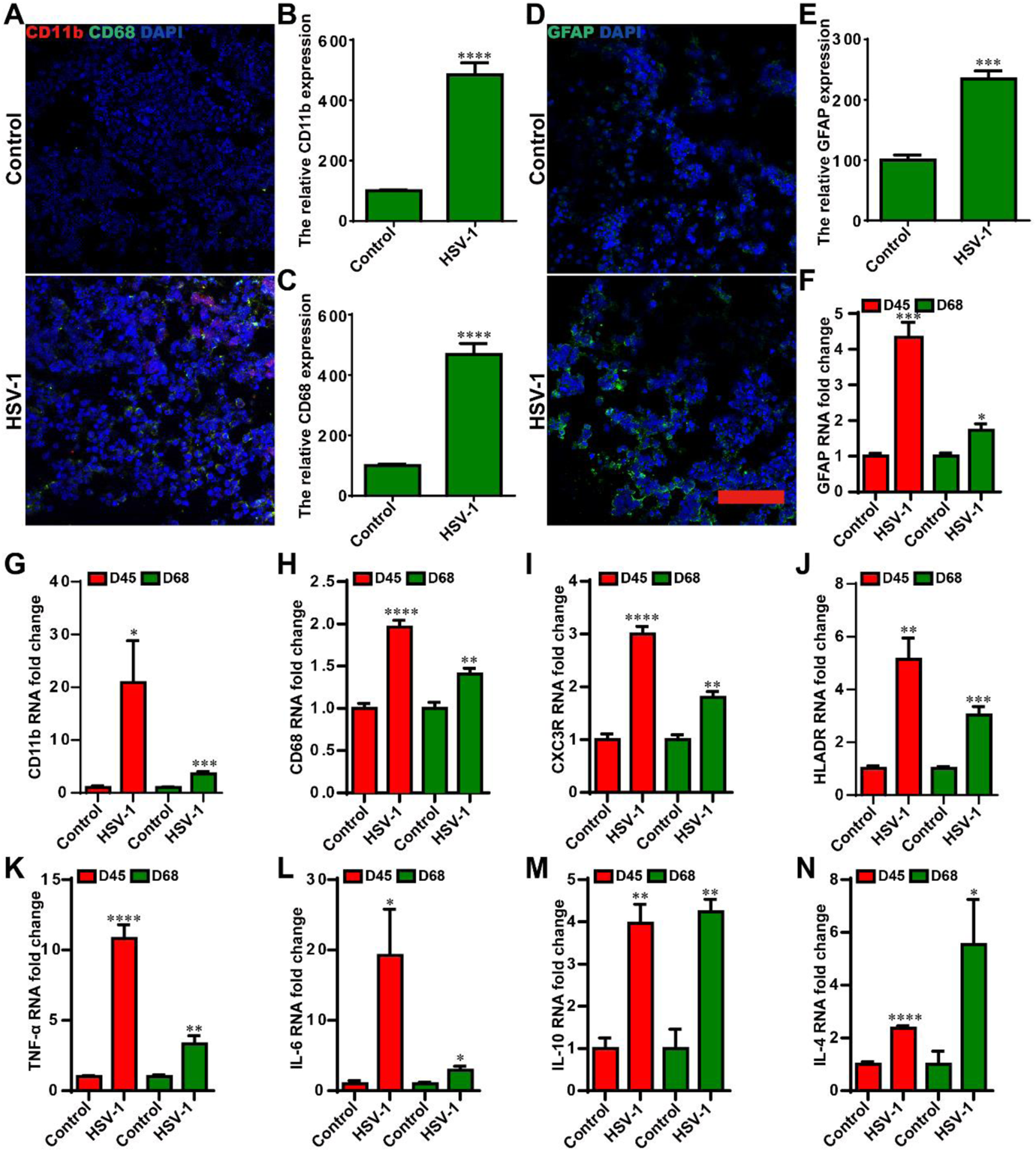
HSV-1 infection led to reactive gliosis and neuroinflammation in the cerebral organoids. (A) Co-immunostaining of CD11b and CD68 were identified by immunofluorescence analysis in the HSV-1 infected cerebral organoids at D65+3. (B and C) The relative fluorescence intensity statistics of CD11b and CD68 expressions as illustrated in (A). (D and E) The expression of GFAP was identified by immunofluorescence staining, and the quantifications for relative fluorescence intensity statistics of GFAP expressions were shown. (F) The mRNA of GFAP was monitored by RT-PCR in the HSV-1 infected cerebral organoids at D42+3 and D65+3. (G-J) Validation by RT-PCR of microglia cells markers (CD11b, CD68, CX3CR, and HLADR) in cerebral organoids at D42+3 and D65+3. (K-N) The mRNA expressions were examined for the inflammatory cytokines (TNF-α, IL-6, IL-10, and IL-4) using RT-PCR in cerebral organoids in the different groups at D42+3 and D65+3. The expression value was normalized to the GAPDH expression level. Data represent the mean ± SEM. *p<0.05 vs the control group, **p<0.01 vs the control group, ***p<0.005 vs the control group, ****p<0.001 vs the control group, #p<0.05 vs the HSV-1 infection group, (n=6).

Next, we examined the neuroinflammatory responses in the cerebral organoids at D42+3 and D65+3. The HSV-1 infection induced increased mRNA expressions of the pro-inflammatory mediators (TNF-α and IL-6) and anti-inflammatory mediators (IL-10 and IL-4) (Fig. 4K to 4N). These observations demonstrated that HSV-1 infection led to reactive gliosis and neuroinflammation.

### RBV and VCV treatments resulted in decreased Aβ deposition and normalized AD-mediator expression in the cerebral organoids

RBV, the synthetic guanosine analog, has been widely used as a broad-spectrum antiviral drug in clinical. In contrast, VCV is one of the most commonly used antiviral agents specifically targeting the HSV DNA replication.

To investigate the effects of these two antiviral drugs on Aβ deposition and AD mediator gene expressions in cerebral organoids at the different growth stage D42 and D65, VCV or RBV was added concurrently with HSV-1 infection for 3 days. The AD mediator genes, including PSEN1, PSEN2 and BACE, have abnormal expressions in the HSV-1 infected-cerebral organoids at D42+3 and D65+3; both RBV and VCV effectively revised the HSV-1-induced changes of AD mediators (Fig. 5A to 5C).

**Fig. 5.**
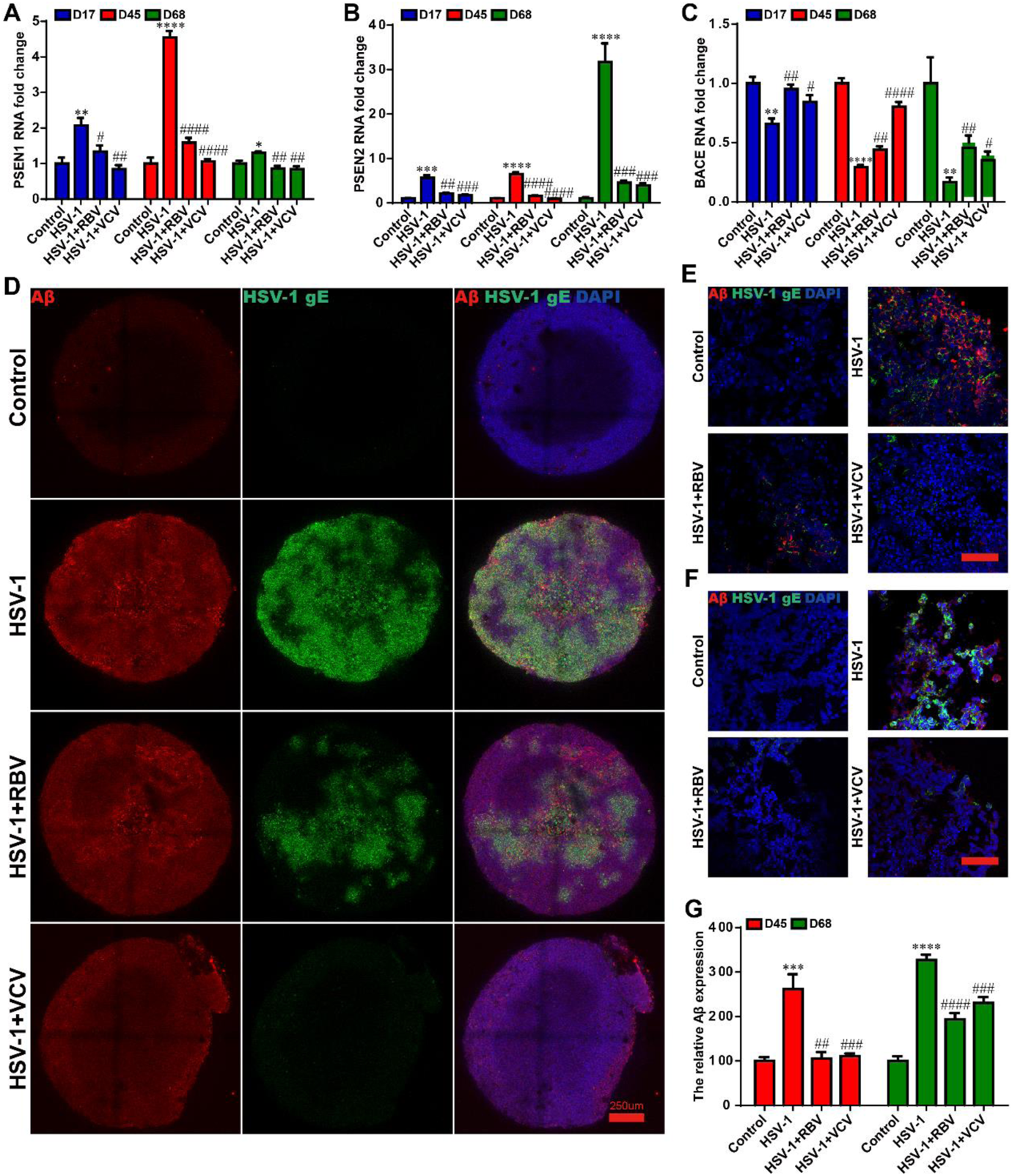
RBV and VCV treatments resulted in decreased Aβ deposition and normalized AD-mediator expression in the cerebral organoids. (A-C) The mRNA expressions of AD mediator genes (PSEN1, PSEN2 and BACE) were examined by RT-PCR in cerebral organoids in different groups at D14+3, D42+3 and D65+3. (D) Whole-mount 3D immunofluorescence imaging of HSV-1 and Aβ in the HSV-1-infected cerebral organoids treated with RBV or VCV at D14+3. Scale bars: 250µm. (E and F) Immunocytochemistry analysis and (G) quantification of the HSV-positive and Aβ-positive cells in the HSV-1 infected cerebral organoids with the RBV or VCV treatment at D42+3 and D65+3. Scale bars: 100µm. Data represent the mean ± SEM. *p<0.05 vs the control group, **p<0.01 vs the control group, ***p<0.005 vs the control group, ****p<0.001 vs the control group, #p<0.05 vs the HSV-1 infection group, ##P<0.01 vs the HSV-1 infection group, ###p<0.005 vs the HSV-1 infection group, ####p<0.001 vs the HSV-1 infection group, (n=6).

To further explore the therapeutic effects of the antiviral drugs on the Aβ deposition. The 3D immunofluorescence analysis showed that the Aβ deposition induced by HSV-1 infection was reduced in the both RBV and VCV groups in the cerebral organoids at D14+3 (Fig. 5D). Concomitantly, we observed a similar reduction in HSV-1 Aβ deposition under the treatment of RBV or VCV in the cerebral organoids at D42+3 and D65+3 compared to the non-treatment group (Fig. 5E and 5F). Quantitative image analysis (Fig. 5G) displayed that the Aβ expression was increased in the HSV-1 infection group, while the Aβ expression was significantly suppressed by both the RBV and the VCV treatments. Together, these observations provide strong preclinical evidence for the therapeutic effects of the RBV and the VCV on the HSV-1-induced Aβ deposition and aberrant AD mediator expressions.

### RBV and VCV treatments reduced reactive gliosis and neuroinflammation in the cerebral organoids

To validate the inhibition of reactive gliosis and anti-neuroinflammation effectivity of these two antiviral drugs in the HSV-1 infected cerebral organoids. We incubated VCV or RBV concurrently with HSV-1 infected cerebral organoids of different stages (D42 and D65) for 3 days. Our results indicated that the expressions of microglia markers (CD11b and CD68) and astrocyte marker GFAP were significantly higher in the HSV-1-infected cerebral organoids compared to the organoids in the control group. Furthermore, the HSV-1-induced overexpression of CD11b, CD68, and GFAP were significantly suppressed in the RBV and the VCV groups by the immunofluorescence staining and quantitative image analysis (Fig. 6A to 6E), compared to those expression in the HSV-1 infection group, suggesting the therapeutic roles of RBV and VCV in the HSV-1-induced gliosis. Concomitantly, we observed that RBV and VCV treatments decreased the expressions of CD11b, CD68, and GFAP at mRNA levels in contrast to the results obtained with HSV-1-infected groups (Fig. 6F to 6H). Moreover, the HSV-1 exposure dramatically increased the mRNA expressions of HLADR and CXC3R, coinciding with increase of microglia markers. And the RBV and the VCV treatment successfully rescued the HSV-1-induced gliosis (Fig. 6I to 6J).Next, we examined the neuroinflammatory responses in the HSV-1 infected cerebral organoids. As shown in Fig. 6K-6N, HSV-1 infection strongly increased mRNA expressions of pro-inflammatory mediators (TNF-α and IL-6) and anti-inflammatory mediators (IL-10 and IL-4), which were evidently turned down by RBV or VCV treatment.

**Fig. 6.**
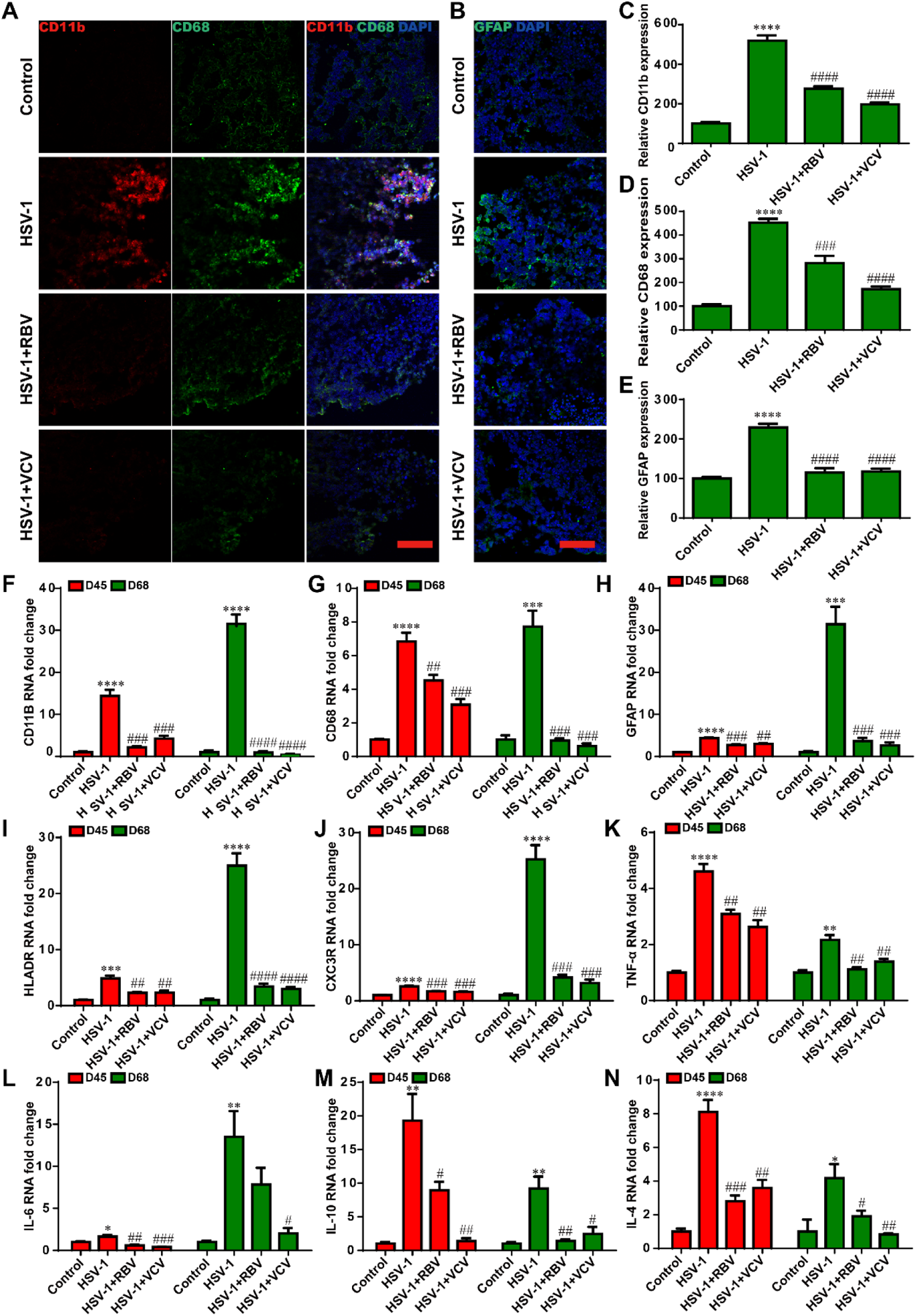
RBV and VCV treatments reduced reactive gliosis and neuroinflammation in the cerebral organoids. (A) Co-immunostaining of CD11b and CD68 were identified by immunofluorescence analysis in the HSV-1 infected cerebral organoids treated with RBV or VCV at D65+3. (B and C) The relative fluorescence intensity statistics of CD11b and CD68 expressions were shown in different groups. (D) The expressions of GFAP were identified by immunofluorescence staining, and the quantifications (E) for relative fluorescence intensity statistics were shown. (F-J) The mRNA expression of microglia cell makers (CD11b, CD68, HLADR and CXC3R) and astrocyte cell marker (GFAP) were examined by RT-PCR in cerebral organoids in the different groups at D42+3 and D65+3. The expression value was normalized to the GAPDH expression level. (K-N) Quantification of the pro-inflammatory mediators (TNF-α and IL-6) and anti-inflammatory mediator (IL-10 and IL-4) mRNA expressions were shown. Data represent the mean ± SEM. *p<0.05 vs the control group, **p<0.01 vs the control group, ***p<0.005 vs the control group, ****p<0.001 vs the control group, #p<0.05 vs the HSV-1 infection group, ##P<0.01 vs the HSV-1 infection group, ###p<0.005 vs the HSV-1 infection group, ####p<0.001 vs the HSV-1 infection group, (n=6).

### RBV and VCV treatments reduced the neuron loss

To further demonstrate the neuroprotective effects of these two antiviral drugs in the HSV-1 infected cerebral organoids. We also investigated the therapeutic effects of RBV and VCV on neuron loss. Strikingly, immunofluorescence analysis revealed that the cerebral organoids displayed a markedly decrease in the expressions of TUJ and MAP2 after HSV-1 infection at D65+3 (Fig. 7A to 7C), and both the RBV and the VCV treatments had the recovery effect on the neuron fate commitment. Similarly, the HSV-1-infected cerebral organoids exhibited a low level of TUJ and MAP2 (Fig. 7D and 7E) mRNA expression, the RBV and the VCV treatments reversed HSV-1-induced the inhibition of neuronal differentiation. These results reveal that RBV or VCV treatments reversed HSV-1-induced neuron loss.

**Fig. 7.**
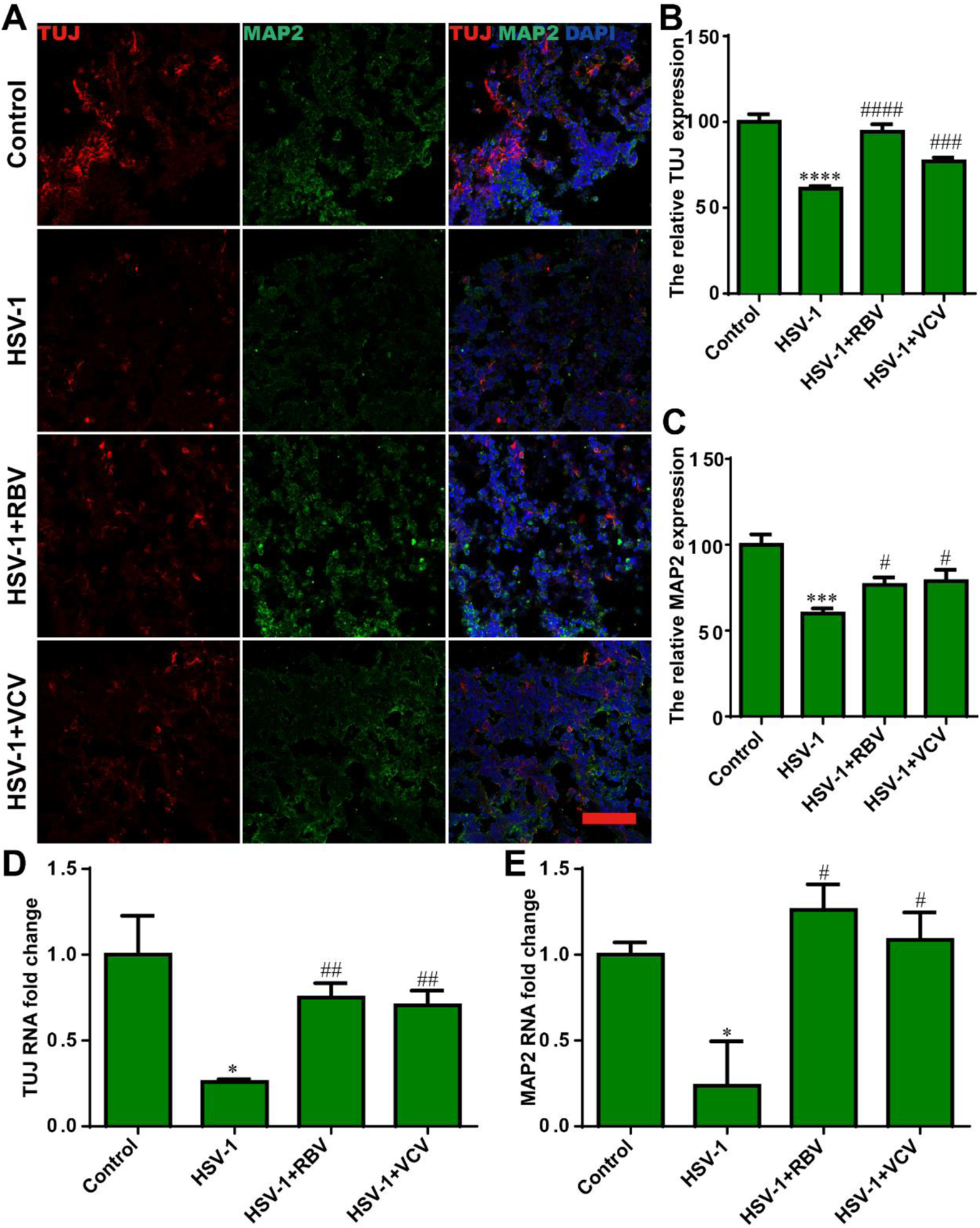
RBV and VCV treatments reduced the neuron loss. (A) Co-immunostaining with TUJ and MAP2 in cerebral organoids in the different group at D65+3. (B and C) The relative fluorescence intensity statistics of CD11b and CD68 expressions in cerebral organoids were shown in different groups at D65+3. (D and E) The TUJ and MAP2 mRNA expressions were quantified by RT-PCR in the cerebral organoids in the different groups at D42+3 and D65+3. Data represent the mean ± SEM. *p<0.05 vs the control group, ***p<0.005 vs the control group, ****p<0.001 vs the control group, #p<0.05 vs the HSV-1 infection group, ##P<0.01 vs the HSV-1 infection group, ###p<0.005 vs the HSV-1 infection group, ####p<0.001 vs the HSV-1 infection group, (n=6).

## DISCUSSION

Recently, there are increasing clinical evidence^25^ and basic researches^26, 27^ indicating that there is strong correlation linking HSV-1 infection with the AD pathogenesis, which supports the AD viral hypothesis.^28^ With the limited access to human brain tissue, the further mechanistic verification of this correlation is highly relying on the AD transgenic mouse models.^11^ However, HSV-1 is a highly human-specific virus and does not infect the wild type mice in the natural settings.^29^ In addition, these transgenic mouse infection models fail to fully capture the human-specific neuropathological features in AD patients due to the distinct differences between humans and mice in terms of polarized neuroepithelium, intricate neurodevelopmental trajectory, and spatiotemporal self-organization in the cerebral cortex. Thus, there is a strong demand to develop more persuasive preclinical models for studying the casual relation between SAD pathogenesis and HSV infection. Recent advance in human cerebral organoids offer a more faithful human-relevant model system to examine the AD viral hypothesis. In this work, we used cerebral organoids to investigate the influence of HSV-1 infection on pathogenesis associated with AD. We found that HSV-1-infected cerebral organoids could model multiscale neuropathology across the genetic, cellular, and tissue levels, including the Aβ deposition, dysregulated AD mediators, reactive gliosis, neuroinflammation, and impaired neuronal differentiation, which were associated AD. Furthermore, we discovered that both RBV and VCV significantly reduced HSV-1 replication and rescued these HSV-1-induced multiscale pathologic phenotypes. Our work intended to provide high-fidelity human brain infection models to evaluate the essential roles of HSV-1 infection on the pathogenesis of AD, with the application in identifying the effects of drug candidates.

The Aβ deposition is a typical hallmark pathology of AD. In our study, we demonstrated that HSV-1 infection resulted in the Aβ deposition in the cerebral organoids, which was reminiscent of amyloid plaques found in AD patients. In general, the Aβ homeostasis of in human brain is determined by the three kinetic factors: (i) the Aβ generation, (ii) the transport of Aβ across the BBB, and (iii) the Aβ degradation in the brain parenchyma.^30^ The AD mediator genes closely regulate these three kinetic factors in the brain. For example, the β-secretase (BACE) and γ-secretase subunits (PSEN1 and PSEN2) are responsible for the Aβ generation.^31^ APOE, the only cholesterol transporter in the brain, participates in both the Aβ transport across the BBB^32^ and the Aβ degradation^30^. Notably, a recent Bordeaux-3C prospective cohort study^33^ assessed the 10-year risk of AD in 1037 participants and found that the APOE4 carriers with frequent reactivations of HSV-1 had a three-fold increased risk of AD, suggesting the interaction between APOE gene and HSV-1 in AD. However, it is not clear whether HSV-1 may influence the AD mediator genes in human brain. In our study, we found the AD mediator genes PSEN1, PSEN2, ASCL, and EPHB in the cerebral organoids were significantly upregulated, meanwhile the BACE expression was decreased in response to the HSV-1 infection. These findings for the first time clarified that the HSV-1 infection influence the AD mediator genes in the cerebral organoids.

The inability to faithfully recapitulate pathophysiological features of human diseases in the traditional cell culture and animal models causes that most preclinically-validated drug candidates failed in the clinical trials. Specifically, in the pipeline of AD drug discovery, 99.6% of AD candidate drugs that showed the promising results in the traditional models failed in clinical trials.^34, 35^ Currently, only five treatment options have been approved to treat the cognitive symptoms of AD, but these treatments have only modest benefit on cognition and no effect on reversing the disease progression.^36, 37^ Here, we discovered that candidate drugs RBV and VCV had the ability to rescue AD-associated multiscale neuropathology in the HSV-1 infected cerebral organoids. VCV, targeting to herpes simplex and herpes zoster, is approved as the antiviral medication for the infections of HSV-1, HSV-2, herpes zoster and chickenpox. Recently, VCV is repurposed as the first-ever anti-AD drug in phase II clinical trial where 130 mild AD patients (MMSE range 20-28) with positive HSV-1 or HSV-2 serum antibodies are undergoing a 18-month, randomized, double-blind, placebo-controlled trial^38^. Furthermore, a recent retrospective cohort study^9^ performed in Taiwan suggested that VCV could attenuate the risk of dementia in patients with HSV infections. In this study, we found that this anti-HSV drug could significantly repair the HSV-1-induced neuropathology associated with AD in the cerebral organoids, including the dysregulated AD mediators, impaired neuronal differentiation, abnormal microglial activation, and neuroinflammation. Together with the retrospective cohort study, we consider VCV is a promising AD candidate drug and recommend to accelerate its clinical trials.

Ribavirin (RBV) is a broad-spectrum antiviral drug^39^ against a wide range of DNA and RNA viruses. RBV has been introduced to treat chronic hepatitis C virus (HCV) infection^40^ for decades. What is more, RBV is repurposed candidate drug for the cancer treatment.^41^ In our result, RBV rescued the AD-related neuropathological phenotypes associated with AD in the cerebral organoids. Our findings strengthen the potency of RBV as a candidate or part of drug combination therapy for the AD treatment.

Recently, emerging human physiologically relevant in-vitro organotypic and histotypic models offer exciting alternative platform for the basic and preclinical researches in the biomedical field. These models bridge the gaps between the single cell level and the organismal level in the traditional biological model systems and potentially provide more accurate understanding for human diseases and faithful evaluation for drug candidates. To establish these models, organoids and microfabricated scaffolds represent two critical but distinct approaches based on the strategies of developmental biology and bioengineering, respectively. Organoids allow generating miniaturized organs from stem cells in vitro by mimicking organogenesis at the human embryonic development stage. In contrast, the microfabricated scaffolds generate faithful tissue-like structures by reverse engineering and bionic fabrication using bioactive materials and living cells. These two models have demonstrated great potential to address the long-lasting challenge in elucidating the ultimate etiology and the underlying mechanism of AD onset and progression. For example, Dana M. Cairns et al.^42^ pioneered a 3D bioengineered brain model by seeding NSCs into the silk protein scaffold based microdevice to study the effects of HSV-1 infection on AD. Using this bioengineered brain model, they found HSV-1 infection led to the formation of AD-like phenotypes, including the Aβ formation, gliosis, neuroinflammation, and diminished neural network functionality. However, this model could not mimic the cortical cellular diversity and spatiotemporal self-organization in the human cerebral cortex. The apparent cortical structure changes and cortical dysfunctions are indeed critically neuropathological features in AD patients.^43, 44^ The cerebral organoids and the bioengineered brain models can perfectly supplement each other with their benefits. The cerebral organoids explore the power of developmental biology to reproduce the different stages of brain development, especially, the spatiotemporal self-organization cerebral cortex. In summary, our study examined the effects of HSV-1 infection on the cerebral organoids and found that the HSV-1 infection cause multiscale neuropathology associated with AD across the multiple bio-hierarchical levels at the different neurodevelopmental stages. Furthermore, we discovered that antiviral drugs RBV and VCV have potential value in the onset and progression of AD. Overall, our study supports the rationality of the AD viral hypothesis and identifies RBV and VCV as promising drug candidates for AD treatments. Further efforts towards building better HSV-1-infected brain models will be to build brain organoid-on-a-chip, which may include blood-brain barrier, multiscale vascular structure, and precise neuro-microenvironment control over organoids on a microfabricated scaffold device by exploring the benefits from both developmental biology and bioengineering. We expect these new models will offer superior high-fidelity to study the viral mechanisms in neurodegenerative diseases, screen potential drug therapies, and aid precision medicine in the future.

## METHODS AND MATERIALS

### Generation of human cerebral organoids

The human cerebral organoids were generated from H9 hESCs using a previously reported protocol with minor modification.^19^ Detailed composition of cerebral organoid differentiation medium was described in the Table 1. Briefly, Accutase (A1110501, Gibco) was used to disassociate the hESCs into single cells. A total number of 2000 cells were then plated into the hanging drop culture plates (InSphero AG) to form the single embryoid body (EB) in the first 24 hours, and then the EBs were transferred to the ultra-low-attachment 96-well plates (7007, Corning) in the EB formation medium (Table 1) with 4 ng/mL bFGF (100-18B, Peprotech) and 50 mM ROCK inhibitor Y27632. The EBs were fed every other day for 6 days then transferred to the low adhesion 24-well plates in the induction media containing DMEM/F12 (11330032, Gibco), 1× N2 supplement (17502048, Gibco), 1%(v/v) non-essential amino acids (NEAA) (11140050, Gibco), 2 mM GlutaMAX (35050038, Gibco), and 1 µg/mL heparin (H3149, Sigma).

**Table 1.** Composition of Cerebral organoid differentiation medium

| Medium Name | Component | Supplier | Identifier | Volume or concentration |
| --- | --- | --- | --- | --- |
| EB formation medium | mTeSR1 | Stem Cell Technologies | 05850 | 50 mL |
|  | bFGF | Peptotech | 100-18B | 4 ng/mL |
|  | ROCK inhibitor Y27632 | Stem Cell Technologies | 72304 | 50 µM |
| Induction medium | DMEM/F12 | Gibco | 11330032 | 50 mL |
|  | N2 supplement (100×) | Invitrogen | 17502048 | 1 × |
|  | Non-Essential Amino Acids (100×) | Gibco | 11140050 | 1 × |
|  | GlutaMAX (100×) | Gibco | 35050061 | 1 × |
|  | Heparin | Sigma | H3149 | 1 µg/mL |
| Neural expansion medium | DMEM/F12 | Gibco | 11330032 | 24 mL |
|  | Neurobasal medium | Gibco | 21103049 | 24 mL |
|  | N2 supplement (100×) | Gibco | 17502048 | 0.5 × |
|  | B27 supplement - vitamin A (50×) | Gibco | 12587010 | 0.5× |
|  | GlutaMAX (100×) | Gibco | 35050061 | 1× |
|  | Non-Essential Amino Acids (100×) | Gibco | 11140050 | 0.5× |
|  | Human insulin | Sigma | I9278-5ML | 2.5 µg/ml |
|  | beta-mercaptoethanol | Merck | 8057400005 | 25 nM |
| Neural maturation medium | DMEM/F12 | Gibco | 11330032 | 24 mL |
|  | neurobasal medium | Gibco | 21103049 | 24 mL |
|  | N2 supplement (100×) | Gibco | 17502048 | 0.5× |
|  | B27 supplement + vitamin A (50×) | Gibco | 17504044 | 0.5× |
|  | GlutaMAX (100×) | Gibco | 35050061 | 1× |
|  | Non-Essential Amino Acids (100×) | Gibco | 11140050 | 0.5× |
|  | Human insulin | Sigma | I9278-5ML | 2.5 µg/ml |
|  | beta-mercaptoethanol | Merck | 8057400005 | 25 nM |

On day 12, the EBs were embedded in the seated Matrigel droplets (356231, Corning), and these droplets were solidified at 37°C. Embedded EBs were subsequently cultured in the neural expansion medium (Table 1) containing 50% (v/v) DMEM/F12, 50% (v/v) neurobasal medium (21103049, Gibco), 0.5× N2 supplement, 0.5× B27 supplement without vitamin A (12587010, Gibco), 2 mM GlutaMAX, 2.5 ng/mL human insulin (I9278-5ML, Sigma), 0.5% (v/v) non-essential amino acids, and 25 nM beta-mercaptoethanol (8057400005, Merck). The embedded EBs were cultured in the stationary condition in 6-well plates for 4 days. Then the plates were transferred to an orbital shaker (HS-25A, MIULAB) rotating continuously at 75 rpm. The culture medium in the dishes was replaced with the neural maturation medium (Table 1) containing 50% (v/v) DMEM/F12, 50% (v/v) neurobasal medium, 0.5× N2 supplement, 0.5× B27 supplement with vitamin A (17504044, Gibco), 2 mM GlutaMAX, 2.5 ng/mL human insulin, 0.5% (v/v) non-essential amino acids, and 25 nM beta-mercaptoethanol.

### HSV-1 infection and phenotypic rescue

HSV-1 strain F stocks were propagated in the Vero cells (African green monkey kidney), with the DMEM supplemented with 2% (v/v) Fetal Bovine Serum (FBS). The propagation was terminated when 90% of the cells appeared with the cytopathic effect. The virus stocks were harvested by repeatedly freezing and thawing the supernatants and the cells for three times, followed by filtering with the 0.22 µm filter and stored in the −80 °C fridge before use. The plaque assay determined the titers of HSV-1 stocks in the Vero cells. Briefly, the Vero cells were cultivated in 6-well plates and incubated with a series of 10-fold virus dilutions for two hours. Then cells were overlaid with 2 mL/well of DMEM (11054020, Gibco) containing 1.2% (v/v) methylcellulose, 10% (v/v) FBS, 1% (v/v) penicillin-streptomycin solution (P0781, Sigma). Plates were incubated at 37 °C with 5% CO^2^ for five days. Cells were fixed with 4% (w/v) paraformaldehyde (PFA) and stained with 0.5% (w/v) crystal violet solution. HSV-1 strain F stocks were maintained at a stock concentration of 10^7^ pfu/mL.

The cerebral organoids at the different developmental stages (D14+3; D42+3; D65+3) were incubated with the titer of HSV-1(100,000 pfu/organoid) in the culture medium for three days. For the mock infections, an equal volume of control culture medium was used. To investigate the effects of VCV and RBV on the pathological phenotypes of cerebral organoids, VCV or RBV was added at a concentration of 200 μM concurrently with HSV-1 infection. After an additional three days’ incubation, the organoids were completely washed with the neural maturation medium.

### Histology and immunofluorescence

Immunofluorescence analysis was conducted as a previous report described ^23^. The cells were fixed with 4% (w/v) PFA for 15 minutes and blocked with donkey serum for one hour. Blocking and permeabilization was performed using 0.3% (v/v) Triton X-100 and 3% (v/v) normal donkey serum in PBS for 20 minutes. Primary antibodies were then incubated at 4 °C overnight. Subsequently, the cells were incubated with the secondary antibodies at the following concentrations: Alexa Fluor 488 Goat anti-Mouse (1:200 dilution, Life Technologies), Alexa Fluor 568 Goat anti-Mouse (1:200 dilution, Life Technologies), Alexa Fluor 488 Donkey anti-Rabbit (1:200 dilution, Life Technologies) and Donkey anti-Rabbit (1:200 dilution, Life Technologies). The nuclei were counterstained with DAPI (62247, Thermo Scientific).

To perform immunofluorescence analysis, cerebral organoids were fixed in 4% PFA and sectioned into 10 µm thick slices with a cryostat (CM1950, Leica). Images were captured via a Leica TCS SP8 STED confocal microscope equipped with a LAS X software. Detailed antibody staining was described in the Table 2. The same laser intensity, detector sensitivity, amplification value, and offset were used for all micrograph acquisitions for quantification of the same protein expression. The expression level of nuclear and cytosolic proteins were calculated by the following formula.

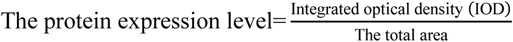

**Table 2.**
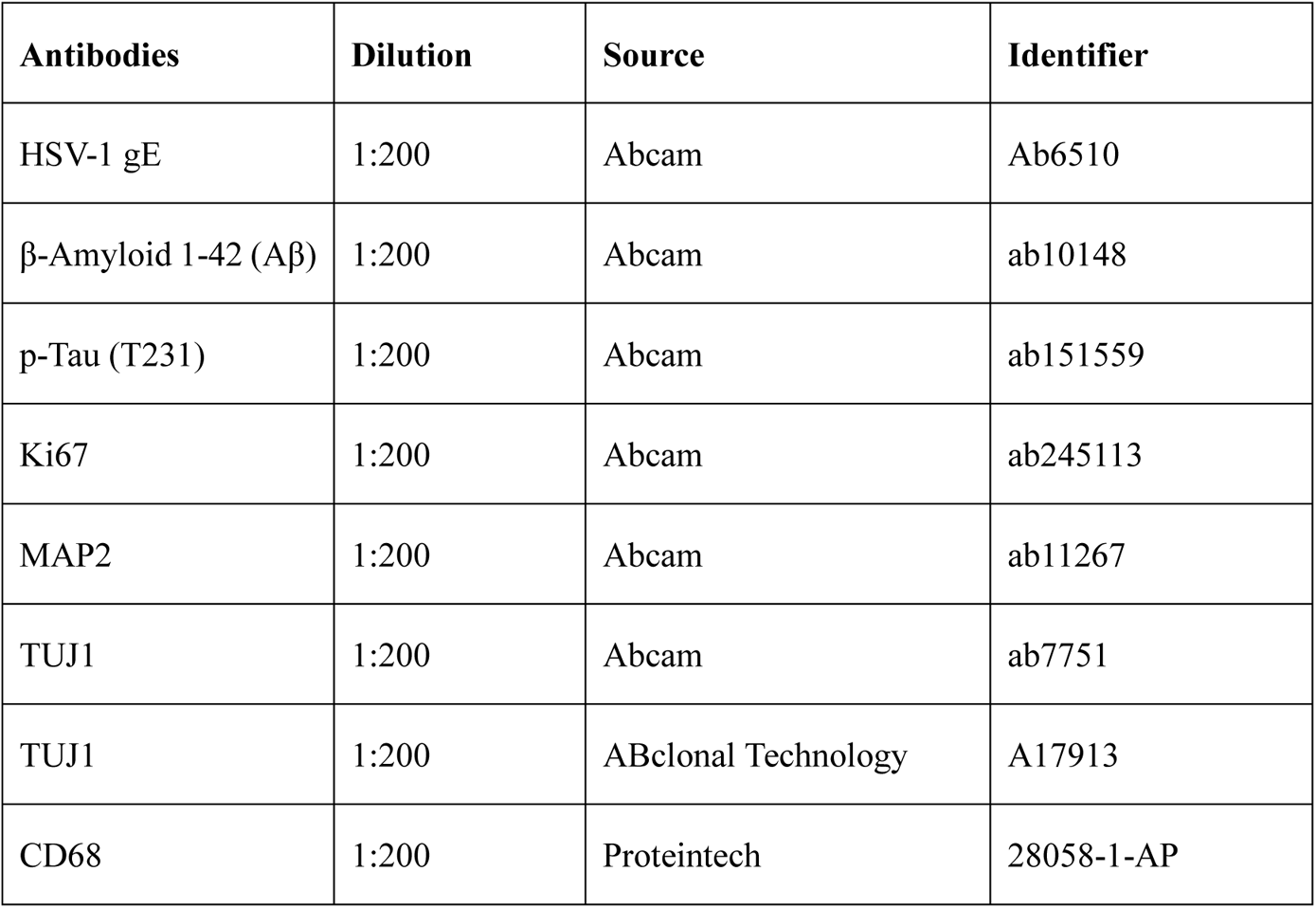
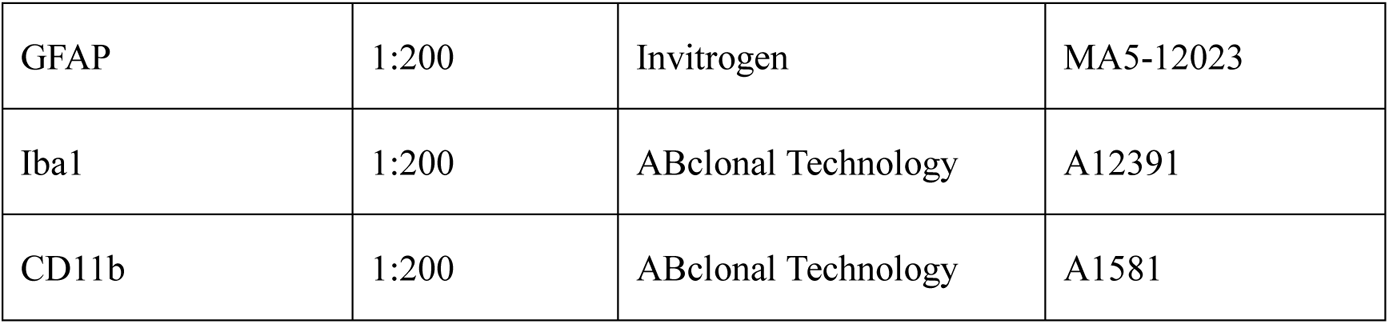
Key resource table

Moreover, each IOD/area value was calculated by subtracting background IOD/area value from the directly measured IOD/area value ^24^. Images analysis was conducted with NIH Image J and LAS X software (NIH, MD, USA).

### Whole-mount Immunostaining of organoids

We performed whole-mount immunostaining followed by confocal microscopy to examine the localizations and organization of the HSV-1-induced Aβ deposition within the cerebral organoids. The organoids were washed with PBS and fixed overnight in 4% (v/v) PFA at 4 °C. After being washed with PBS for five hours, the organoids were blocked overnight at RT in 0.5% (v/v) BSA and 0.125% (v/v) Triton X-100 in PBS. The organoids were incubated in the primary antibodies (anti-HSV-1 gE and anti-Aβ) diluted in 0.5% (v/v) BSA and 0.125% (v/v) Triton-100 in PBS for two days at 4 °C. Unbound antibodies were removed via multiple washes with PBS for one day at RT. Then, the cerebral organoids were then incubated with Alexa Fluor 568 Goat anti-Mouse (1:200 dilution, Life Technologies), Alexa Fluor 488 Donkey anti-Rabbit (1:200 dilution, Life Technologies) and Donkey anti-Rabbit (1:200 dilution, Life Technologies) for four hours following nuclei staining with DAPI (1:1000 dilution) for two hours. 3D image stacks (100 µm in the thickness) were acquired for representative organoids. The interval between the neighboring stacks was 4 µm.

### Real-time PCR

Total mRNAs were isolated from the cerebral organoids or the cells using Trizol, then cDNA was synthesized using ABScript III RT Master Mix (RM21452, ABclonal Technology). RT-PCR was performed using SYBR Green Real-time PCR Master Mix (RK21203, ABclonal Technology) under the following reaction conditions (35 cycles): denaturation at 95 °C for 1 minute, annealing at 58 °C for 30 seconds, and extension at 72 °C for 30 seconds. Primer sequences were described in Table 3. The expression levels were normalized relative to the expression of the housekeeping gene GAPDH using the comparative Ct–method 2^−ΔΔCt^.

**Table 3.**
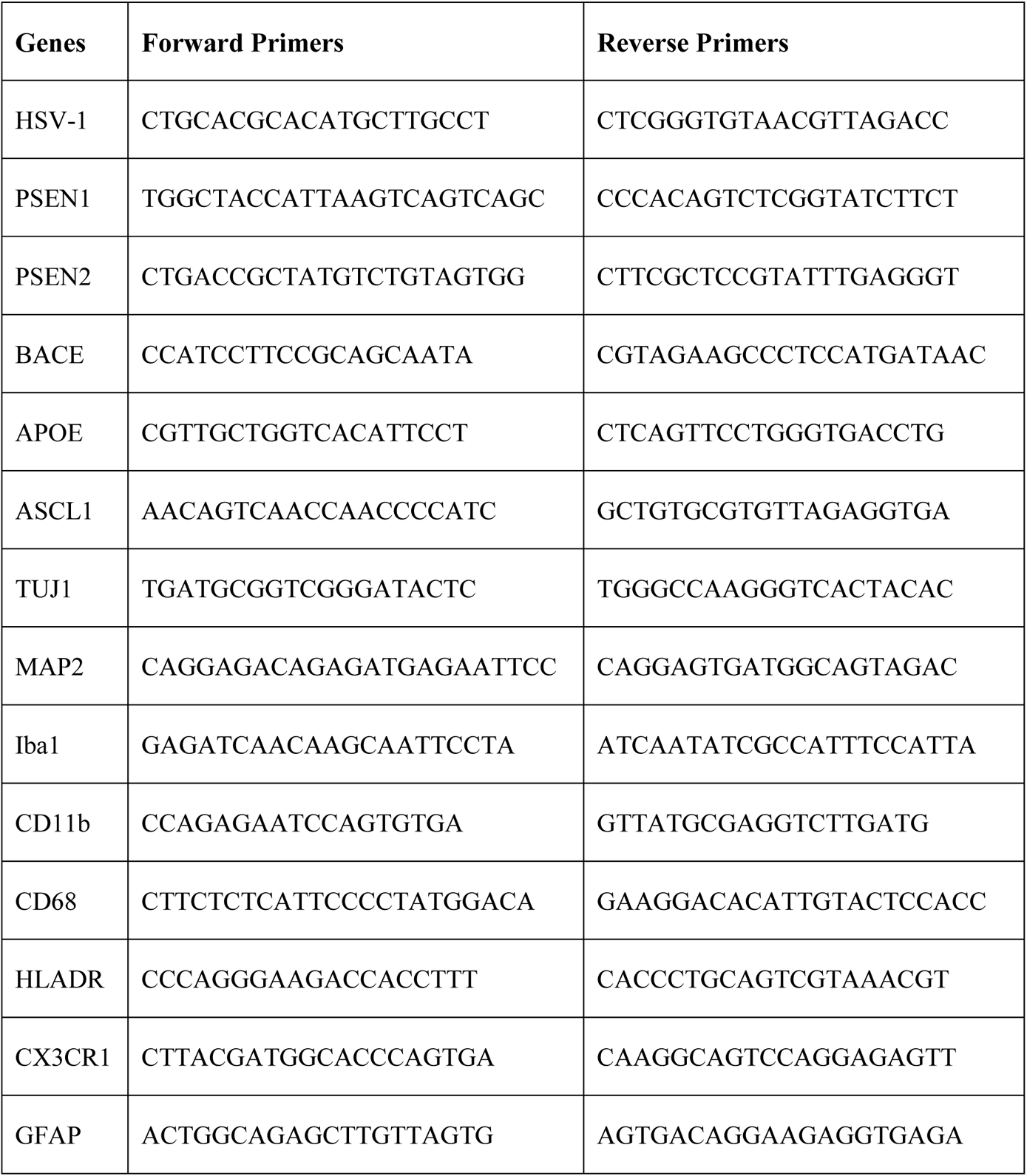
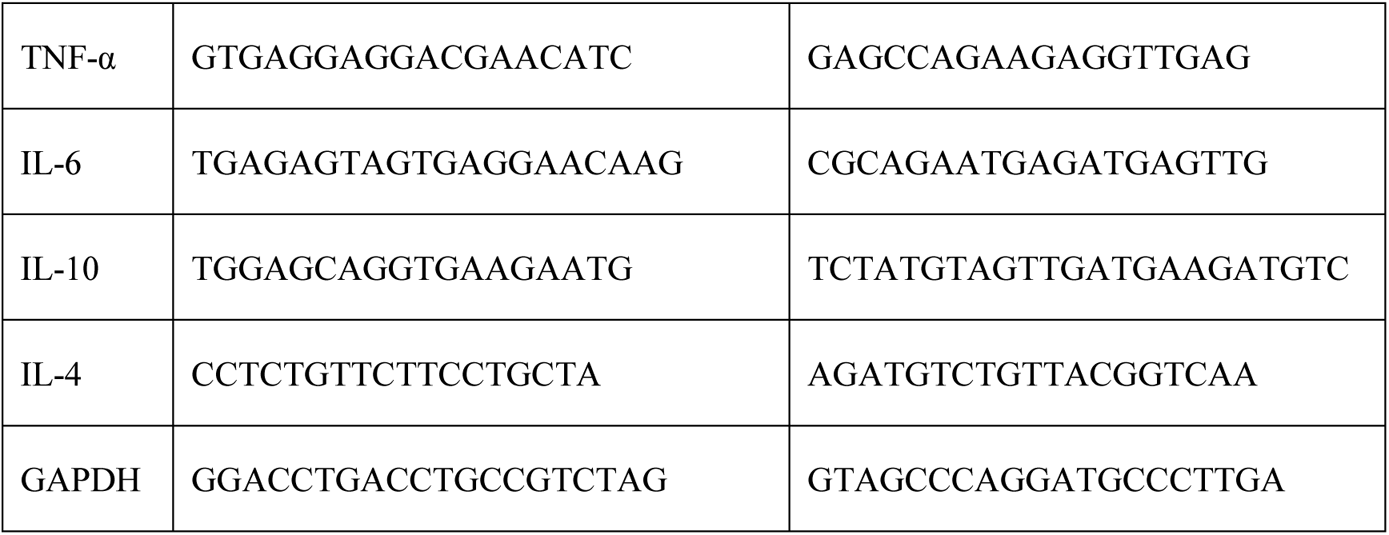
Primer sequences used for RT-PCR

### RNA-Seq and Data Analysis

The PSC-derived cerebral organoids were infected with the HSV-1(1:100,000 pfu/organoid). The inocula were removed after three days of infection. The cerebral organoids were harvested, and the RNAs were extracted. RNA sequencing was performed using an Illumina NextSeq 6000 with an average of 20 million reads per run. The Gene Ontology (GO) enrichment analysis, Kyoto Encyclopedia of Genes and Genomes (KEGG) pathway enrichment analysis were performed using DAVID database (https://david.ncifcrf.gov/). In order to get astringent DEG data set, only DEG with >= 1.5 fold change was used for GO enrichment analysis and KEGG pathway enrichment analysis.

### Statistical analysis

Statistical analysis of data was expressed as means ± SEM. Student’s t-tests were applied to data with equal or more than two groups. ANOVA analysis was used for comparing data with greater than two groups. In all the analyses, group differences were considered statistically significant as follows: ∗p< 0.05, ∗∗p< 0.01, ∗∗∗p< 0.005, ∗∗∗∗p< 0.001. Sample sizes were indicated in the figure legends.

## ACKNOWLEDGEMENTS

This study was supported in part by grants from the National Natural Science Foundation of China (Grant No. 31871018, http://www.nsfc.gov.cn/), received by P.C. The National Natural Science Foundation of China (Grant No. 32122008, http://www.nsfc.gov.cn/) and the National key Research and Development Plan of China (Grant No. 2021YFC2300200), received by Y.W. The funders had no role in study design, data collection and analysis. Thanks for technical support from Research Center for Medicine and Structural Biology of Wuhan University, and Innovations in stem cell and organoids project (ISCO).

## AUTHOR CONTRIBUTIONS

The author contributions are listed according to Contributor Roles Taxonomy (CRediT).

Conceptualization: Pu Chen.

Data curation: Haowen Qiao, Wen Zhao.

Formal analysis: Haowen Qiao, Wen Zhao, Pu Chen.

Funding acquisition: Pu Chen, Ying Wu.

Investigation: Haowen Qiao, Wen Zhao, Pu Chen, Moujian Guo, Tao Chen, Lili Zhu, Ying Wu.

Methodology: Haowen Qiao, Wen Zhao, Pu Chen.

Project administration: Pu Chen.

Resources: Pu Chen, Ying Wu.

Software: Haowen Qiao, Wen Zhao, Jibo Wang.

Supervision: Pu Chen, Ying Wu, Zhentao Zhang.

Validation: Wen Zhao, Pu Chen.

Visualization: Haowen Qiao, Wen Zhao, Jibo Wang.

Writing – original draft: Haowen Qiao, Pu Chen.

Writing – review & editing: Pu Chen, Ying Wu, Zhentao Zhang.

## CONFLICT OF INTEREST

P.C. is a founder of, and has an equity interest in: (i) Shenzhen Convergence Bio-Manufacturing Co., Ltd., a company that is developing convergence bio-manufacturing technologies to enable regenerative medicine, and (ii) Organoid chip Medical Co.,Ltd., a company that is developing organoid-on-a-chip technologies for clinical solutions. P.C.’s interests were viewed and managed in accordance with the conflict of interest policies.

## DATA AVAILABILITY STATEMENT

All data that support the findings of this study are included within the article.

